# Local structure of DNA toroids reveals curvature-dependent intermolecular forces

**DOI:** 10.1101/2020.07.23.211979

**Authors:** Luca Barberi, Françoise Livolant, Amélie Leforestier, Martin Lenz

## Abstract

In viruses and cells, DNA is closely packed and tightly curved thanks to polyvalent cations inducing an effective attraction between its negatively charged filaments. Our understanding of this effective attraction remains very incomplete, partly because experimental data is limited to bulk measurements on large samples of mostly uncurved DNA helices. Here we use cryo electron microscopy to shed light on the interaction between highly curved helices. We find that the spacing between DNA helices in spermine-induced DNA toroidal condensates depends on their location within the torus, consistent with a mathematical model based on the competition between electrostatic interactions and the bending rigidity of DNA. We use our model to infer the characteristics of the interaction potential, and find that its equilibrium spacing strongly depends on the curvature of the filaments. In addition, the interaction is much softer than previously reported in bulk samples using different salt conditions. Beyond viruses and cells, our characterization of the interactions governing DNA-based dense structures could help develop robust designs in DNA nanotechnologies.

## INTRODUCTION

DNA is a negatively charged semi-flexible polymer, which causes it to form extended coils when placed in water. It however adopts a very different conformation *in vivo*, where it is tightly packaged and segregated within the prokaryotic cytoplasm or the eukaryotic nucleus. Even more extreme confinements are achieved in sperm cells and certain viral capsids, such as bacteriophages, where physical contact between DNA strands becomes a major constraint on the polymer’s conformation [1]. Similar densities and/or curvatures are achieved in man-made objects such as toroidal DNA bundles [2, 3] and DNA origami [4–6] designed for applications as diverse as gene and drug delivery [7, 8] membrane deformation and permeabilization [6, 9], or the design of chiral plasmonic metamolecules [10].

In these dense packings, the center-to-center distances between DNA helices can be as small as 2.5 to 3 nm. Such strong confinements require overcoming the electrostatic repulsions between DNA strands, which can be achieved through a variety of mechanisms such as ATP-dependent compaction in bacteriophage chromosomes [1, 11], Watson-Crick base-pairing in origami folds [12, 13], macromolecular crowding [2, 14] or cation-induced condensation [14–16], which plays a major role. The electrostatic forces between helices may turn from repulsive to attractive thanks to the presence of high-valence cations such as spermine (4+) and spermidine (3+) or small proteins such as protamines and H1 histones [17–20]. Several theoretical accounts of these effects have been proposed, suggesting that the hydration forces [21–23], ion chemisorption [24], structuring of the ion clouds [25, 26] or bridging [27–29] may all play a role in the effective DNA-DNA attraction, although no consensus exists as to their relative contributions.

Experimental characterizations of these complex interactions have largely relied on osmotic compression, whereby an ordered array of condensed DNA helices is equilibrated against a salt solution also containing a large sterically excluded polymer [21]. These experiments have revealed that the interactions between aligned chains are well approximated by the sum of a short-ranged exponential repulsion that is relatively insensitive on the DNA’s ionic environment and a longer-ranged exponential attraction that strongly depends on it. Single-molecule experiments have also been conducted, yielding some smaller-scale information on the condensation energy induced by these interactions, although not in a controlled geometry [30].

In this paper, we provide a characterization of these interactions under geometrical conditions that better mimic the tight confinement and high curvature encountered *in vivo*. We use the phage/receptor system, a versatile experimental model that can be used to investigate DNA conformations. We specifically investigate toroidal DNA arrays formed upon DNA ejection from phages in the presence of condensing amounts of spermine [31–33], as they provide a wide range of controlled curvatures whose influence on the local DNA structure we monitor with cryo electron microscopy (cryo-EM).

Our observations reveal that within an individual DNA toroid, the local spacing between neighboring helices increases with increasing local curvature. This spacing is further increased in small, highly curved toroids confined inside a viral capsid compared to giant unconfined toroids. We interpret these results using a mathematical model where the mechanical equilibrium of the toroidal assembly is given by the competition between DNA bending rigidity and effective inter-helical forces. While this model accounts for the variation of the DNA-DNA spacing within a toroid, it also predicts that small confined toroids should have a smaller spacing than giant confined toroids, in contradiction with our observations. This contradiction implies that interactions between highly curved DNA filaments are quantitatively different than at lower curvatures, an effect which we directly quantify by combining our model and our experimental data on the local inter-helical spacing. We also report a much softer interaction between curved DNA helices compared to previous studies on straight DNA arrays, which further suggests that helix-helix interactions depend on curvature.

Overall, our results highlight the role of the competition between curvature-dependent inter-helix interactions and the elastic response of single filaments in shaping dense DNA assemblies.

## RESULTS

### We use DNA ejection from phages to produce small confined DNA toroids and giant unconfined DNA toroids

The phage/receptor system has previously been used to produce DNA toroids, from small monomolecular toroids consisting of a portion of the phage genome confined within its capsid [32] to giant assemblies of multiple DNA molecules ejected from many phages [31]. Inspired by these works, we produce toroids of variable curvature using two phage species (T5 and Lambda) with different capsid sizes. We trigger DNA ejection in the presence of spermine to condense DNA and form toroids. In some experiments, we add DNAse at different times corresponding to different stages of the DNA ejection process. DNAse degrades the already-ejected fraction of the DNA molecule outside the capsid and interrupts the transfer. The remaining DNA is condensed into a toroid confined inside the capsid. In other experiments where DNAse is absent, the genome is fully transferred out [34, 35] and the molecules from several phages coalesce into giant unconfined toroids typically wrapped around one of the capsids.

Cryo-EM examination reveals the variety of confined toroids thus generated inside phage capsids [Fig. 1 (C, D)], as well as unconfined toroids generated outside of them [Fig. 1 (A, B)]. Small toroids formed in the presence of DNAse have their outer radius *R*_out_ fixed by the species-dependent capsid size, namely 36 − 37 nm in phage T5 (Fig. 1C) and 26 nm in phage Lambda (Fig. 1D). Inner radii vary from 7 to 20 nm in both species, and are on average slightly smaller in Lambda. The inner radius of small toroids is unconstrained. Conversely, in giant toroids, the capsid fixes the inner radius 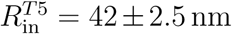, and 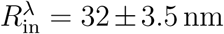, while the outer radius is unconstrained. Giant toroids are relatively monodisperse in all conditions, with outer radii *R*_out_ = 144 ± 15 nm, with no significant difference between toroids obtained from phages T5 and Lambda.

**Figure 1.**
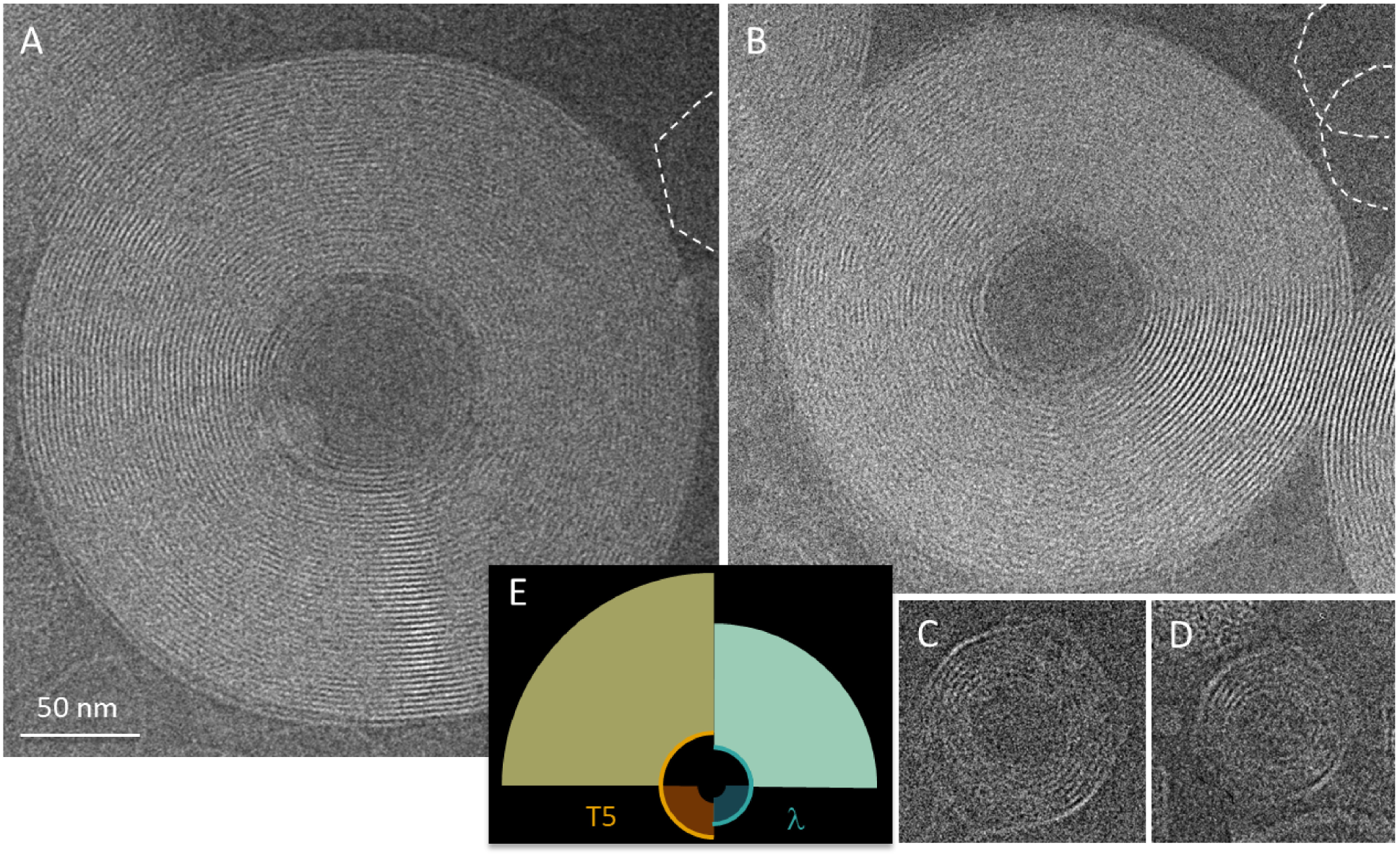
CryoEM imaging of the different types of toroids obtained by DNA ejection from phages in the presence of condensing amounts of spermine. Toroids are here observed in top views. (A, B) Giant unconfined toroids formed outside phage capsids after the ejection of multiple DNA molecules from phage T5 (A) or Lambda (B). Empty capsids are visible in the periphery of the image and are outlined by white dotted contours. *C*_spermine_ = 40 mM. In (A), the ejection is not complete for the capsid located at the core of the giant toroid and the remaining segment inside is condensed into a small confined toroid. (C, D) Small confined toroids trapped in after partial DNA ejection in the presence of DNAse in the external medium, for phage T5 (C) and Lambda (D). *C*_spermine_ = 4 mM. (E) Sketch of the different toroid populations: giant external toroids with fixed inner boundary and small confined ones with fixed outer boundary. In all cases, the radius of the fixed boundary is dictated by the radius of the phage capsid, *i*.*e*. the phage species: here T5 (orange) and Lambda (cyan).

The wide range of well defined toroid sizes generated by our approach allows us to access curved DNA configurations with radii of curvature varying continuously from 7 to 375 nm, as sketched in Fig. 1E.

### Inter-helix spacing is larger in highly curved regions of either types of toroids

Both global and local measurements of inter-helix spacing are possible from the analysis of cryo-electron micrographs. As already shown by other cryo-EM studies of DNA toroids [32, 36], toroids observed in top views exhibit concentric striated patterns (Fig. 2A). The local hexagonal packing of DNA is visualized in side views (Fig. 2C). Wherever we find striations in top views (Fig. 2C’), we observe the hexagonal structure along the direction of one of its lattice planes. The striation spacing is 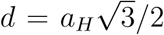, where *a*_*H*_ is the interhelix spacing between DNA segments, as sketched in Fig. 2C”. Note that we do not observe striations along the whole circumference of the toroid (Fig. 2A, asterisks) due to the rotation of the hexagonal lattice in this direction [32] and other defects [36].

**Figure 2.**
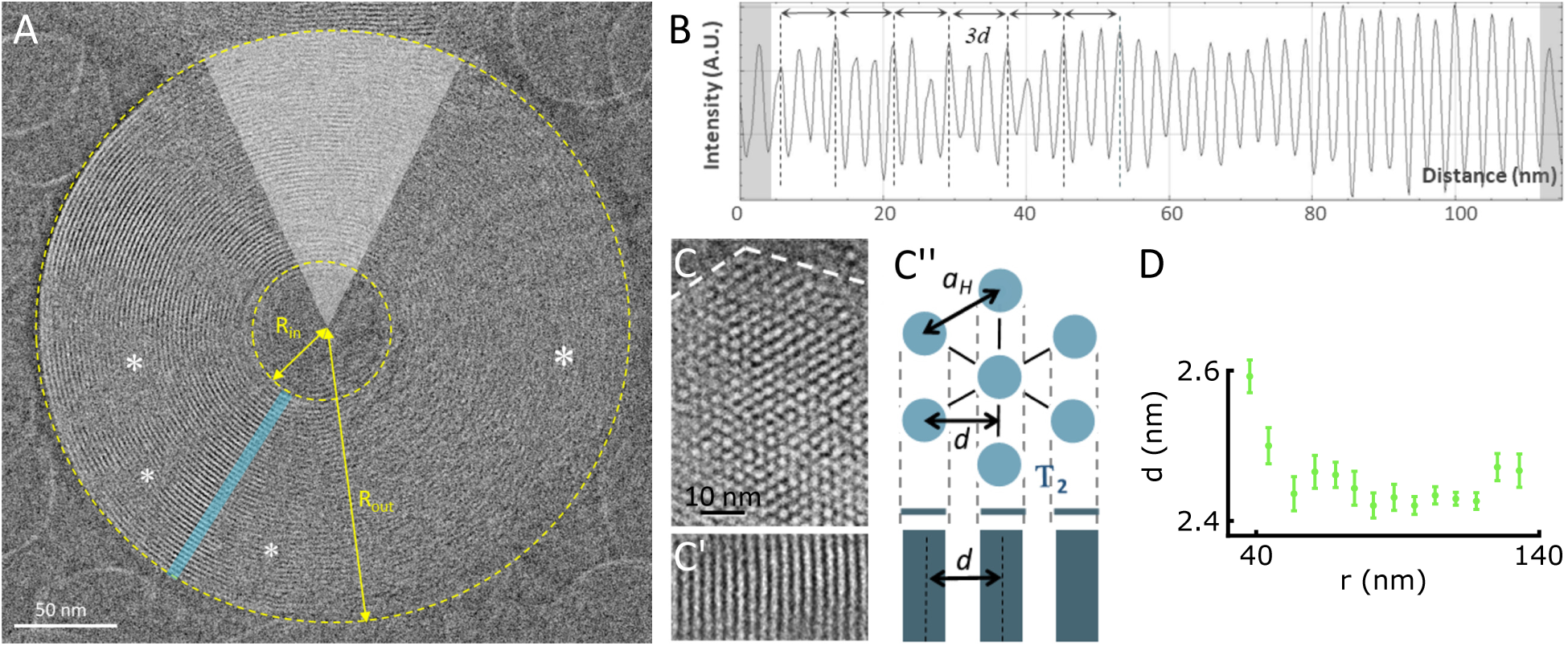
Filament spacing measurements on cryo-EM images (here giant toroids). (A) Top view of a toroid. The inner and outer radii *R*_in_ and *R*_out_ are outlined by yellow dotted lines. Asterisks highlight some regions devoid of striations. An example of a line profile used for our analysis is given in cyan. The pale gray overlay highlights a sector whose outer boundary is constrained by a contact with another toroid, and is therefore excluded from our analysis. (B) Plot of the line profile shown in (A). The striation spacing *d* is measured from averages over 3 layers. The inner and outer layers (gray overlay) are discarded. (C) Side view of a toroid showing the arrangement of its DNA filaments into a triangular lattice in its cross-section. (C’) Detail of the striations observed in a top view projection along a T2 projection axis. (C”) Corresponding sketch of the hexagonal lattice with its projection along the T2 axis into parallel stripes. *d* is the striation spacing and *a*_*H*_ the interhelix spacing. (D) Variation of *d* as a function of *r*, averaged over 10 line profiles recorded on the toroid shown in (A).

To measure the average striation spacing 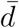, we record line profiles on top views of toroids (Fig. 2A, cyan line) and divide the profile length by the number of layers (Fig. 2B). We find 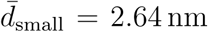 in small toroids and 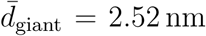 in giant ones (Supplementary Data I). This suggests an increase of 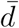 with increasing curvature. The small difference in radii between T5 and Lambda capsids does not lead to any noticeable difference between the two species. Increasing the spermine concentration from 4 to 40 mM does not significantly affect the striation spacing (Supplementary Data I). A similar independence on spermine concentration was previously observed in dense packings of short DNA fragments in the presence of (relatively) high amounts of monovalent cations (≥ 50 mM) [37]. In the following we thus pool the data obtained under these two spermine concentrations together, and conduct a detailed analysis of seven small toroids (five at 4 mM spermine in T5 and Lambda, two at 40 mM spermine in T5) and seven giant ones (six at 40 mM spermine, T5 and Lambda; one at 4 mM, T5) (Supplementary Data II).

To access local variations of the spacing *d* beyond the mean estimate 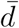, we measure peak-to-peak distances on line profiles. To reduce noise, we take advantage of the large number of layers in giant toroids and average spacings over three layers. On account of the small number of layers in small toroids, the spacings are not averaged there, resulting in noisier data. In all cases, we exclude from the measurements the DNA layers constituting the inner and outer boundaries of the toroids (innermost and outermost layers, shadowed in grey in Fig. 2B) to eliminate the influence of the peripheral fluctuations due to the reduced number of neighboring filaments [38]. In giant toroids, we do not collect data from regions where the toroid is potentially perturbed by a contact with a neighbor (Fig. 2A, pale grey overlay). For each toroid, we record several line profiles (from 8 to 13 in giant toroids, 3 to 6 in small confined ones) and average *d* values. We plot the resulting *d* values as a function of the radius *r* (Fig. 2D, 3).

The result of these local measurements is reminiscent of the trend observed for the mean estimates 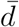, indicating an increase in spacing with increasing curvature. The analysis of giant toroids moreover suggests that this dependence is only relevant at high enough curvatures, as *d* appears roughly constant for curvature radii *r* larger than 75 nm.

### A mathematical model based on the helices’ bending energy accounts for the observed variable spacing

To understand the local correlation between inter-helical spacing and curvature, we develop a mathematical model based on the competition between inter-helical electrostatic interactions and DNA bending rigidity. Qualitatively, imposing curvature on DNA has a high bending energy cost. To relieve this cost, DNA tends to move away from the highly curved inner regions of the toroid to regions with milder curvatures. Due to this tendency to move, we expect DNA to be more closely packed on the outside of the toroid, as observed experimentally. In the presence of strong inter-helical interactions imposing a well-defined value of the inter-helical spacing, these variations tend to be small. Conversely, a relatively large variability in the packing density indicates a loose inter-filament interaction. Our model harnesses this correspondence to infer the inter-filament interaction strength from the spatial variation of their spacing.

We consider a simplified two-dimensional model comprised of concentric DNA circles, reminiscent of the striations appearing in toroid top views (Fig. 2A). The smallest (largest) circle has a radius *R*_in_ (*R*_out_). Denoting the capsid radius by *R*_*c*_, we have *R*_out_ = *R*_*c*_ in small confined toroids while *R*_in_ = *R*_*c*_ in giant unconfined toroids. We model DNA as a semi-flexible polymer, implying that the bending energy per unit length of a filament reads

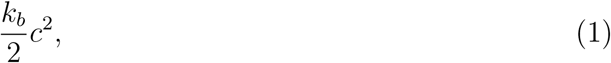

where *c* denotes the local curvature of the filament and where the filament bending stiffness *k*_*b*_ = *k*_*B*_*T*𝓁_*p*_ is related to the persistence length 𝓁_*p*_ of DNA through the thermal energy *k*_*B*_*T*. If DNA were devoid of bending rigidity (*i*.*e*., *k*_*b*_ = 0), the balance between attraction and repulsion between neighboring DNA filaments would fix their spacing to some constant value, *d*_0_, throughout the assembly. This would correspond to a (local) minimum of the interaction potential between neighboring helices. Expanding the interaction energy to second order around this minimum, we write the interaction energy per unit length of a pair of filaments as

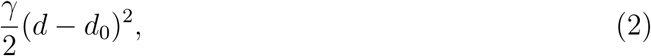

where *γ* is a spring-constant-like, a priori unknown parameter characterizing the strength of the interaction.

Since the typical inter-helical spacing in our set of circles (2.5 − 3 nm) is much smaller than the toroid size, we take the continuum limit of the set of circles and assimilate it to a hollow disk parametrized by the radial coordinate *r* ∈ [*R*_in_, *R*_out_] (Fig. 4). We use the state where the DNA filaments are uniformly spaced by *d* = *d*_0_ as our *reference state*, where the number of circles between *r* and *r* + *dr* is *dr/d*_0_. When *k*_*b*_ > 0, DNA filaments resist bending associated with this state by relaxing local curvature such that the local radius of curvature goes from *r* to *r* + *U* (*r*), where the displacement *U* (*r*) characterizes the magnitude of the deviation of the filament positions from the reference state. In this *final state*, the spacing depends on the position *r* and reads *d*(*r*) = *d*_0_[1 + *U* ^*1*^(*r*)] (Fig. 4), where the prime denotes differentiation with respect to *r*.

**Figure 3.**
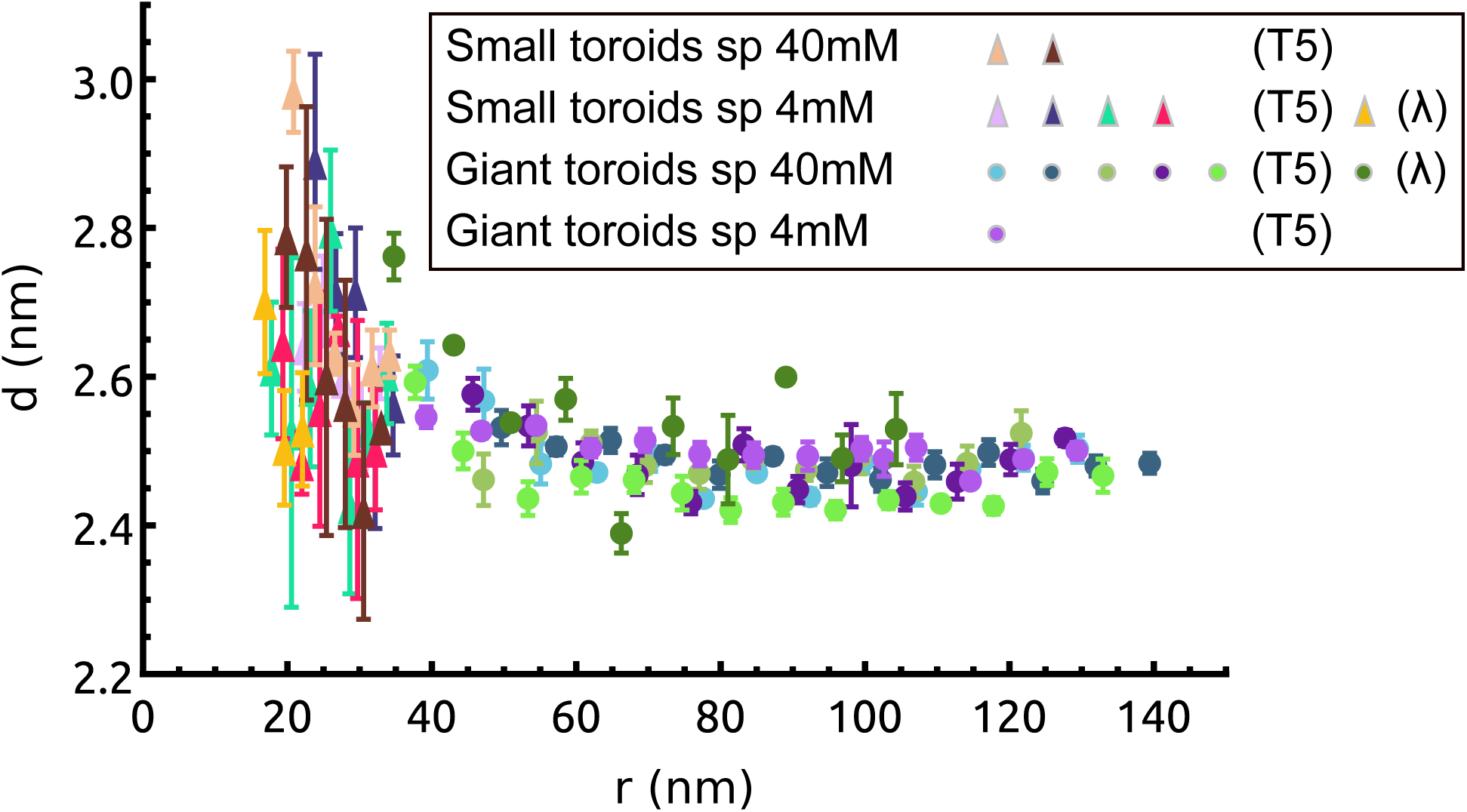
The spacing *d* decreases with increasing distance from the toroid center *r* under all conditions examined. However the curves do not exactly overlap in this representation, hinting at a nontrivial dependence on the toroid radii and boundary conditions which we unravel in our mathematical model.

**Figure 4.**
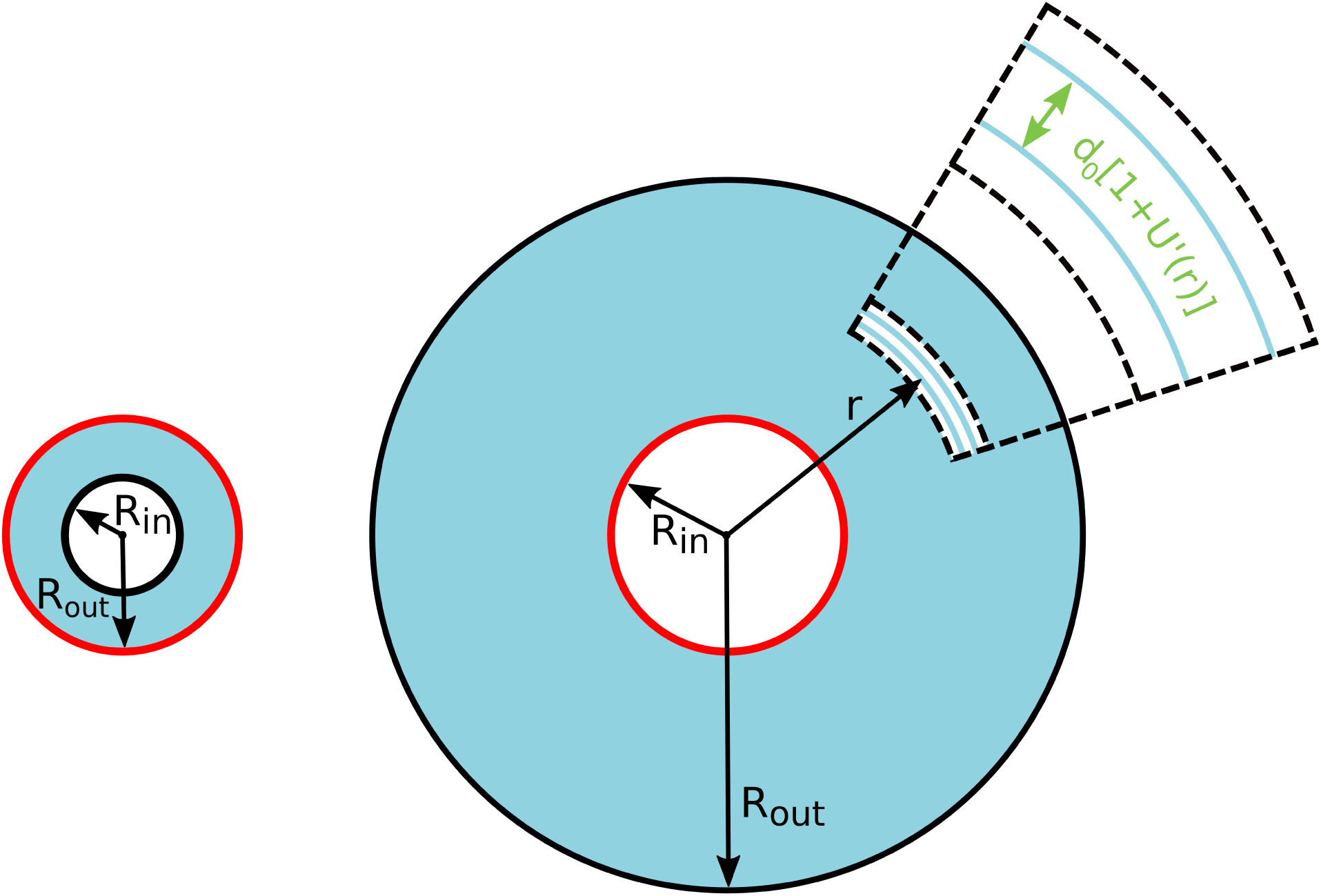
Two-dimensional model of small confined (left) and giant unconfined (right) DNA toroid. Fixed boundaries are colored in red. Both models represent continuum limits for a dense set of concentric circles (cyan). Dashed box: close-up showing two circles, located at *r* and *r* + *dr* in the *reference state* and spaced by *d*(*r*) = *d*_0_[1 + *U* ′(*r*)] in the *final state*.

The relatively small deviations of our filament spacings from a homogeneous distribution (of the order of 10% – see Fig. 3) may be expressed as *U* ′(*r*) *≪* 1, and suggests that bending effects are weak compared to interactions. We formalize this observation by introducing a dimensionless small parameter 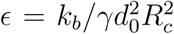 that quantifies the magnitude of the DNA bending stiffness relative to the inter-filament interactions. We assume *ϵ* ≪ 1 and write *U* (*r*) = *ϵu*(*r*) in the following. The two energetic contributions detailed in Eq. (1) and Eq. (2) thus yield a continuum free energy

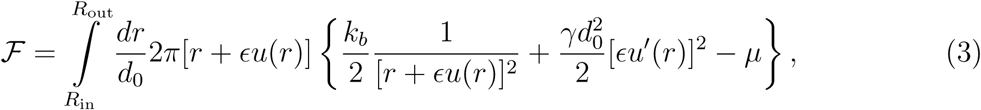

where the third term in the curly brackets features the tension of the DNA filaments, which we introduce as a Lagrange multiplier ensuring that the total DNA length ℒ is conserved:

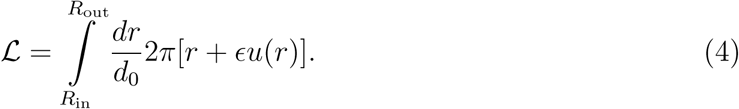

We compute the equilibrium filament displacement *u*(*r*) by minimizing the free energy functional [Eq. (3)] (see Supplementary Mathematical Modelling). To leading order in *ϵ*, we obtain for small toroids

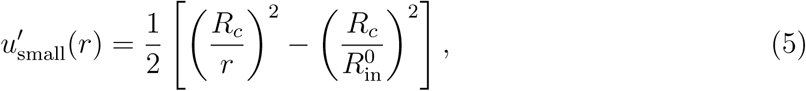

where 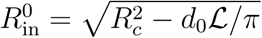. On the other hand, giant toroids yield

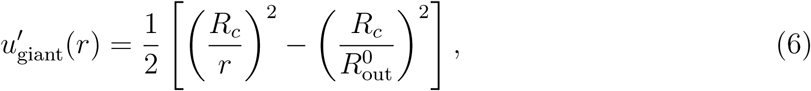

where 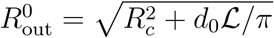.

Thus in both small and giant toroids, our model predicts that the spacing *d* decreases with increasing distance *r* from the center. While this finding is consistent with the data presented in Fig. 3, another prediction of our model appears at odds with our experimental observations. Indeed, our model predicts 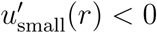, implying 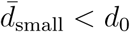 in small toroids, while 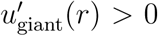, implying 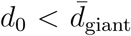 in giant ones. By contrast, our experimental data show 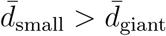. These statements can only be reconciled by assuming that the optimal pacing *d*_0_ has a different value for small vs. giant toroids.

### Fitting of the model’s parameters reveals differences in interactions in small and giant tori

To understand the effect of confinement-induced high curvatures on DNA-DNA interactions, here we extract the values of the optimal spacings 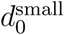 and 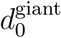 as well as the interaction parameter *γ* from the data presented in Fig. 3. Rather than fitting the individual *d* = *d*(*r*) curves, we exploit our model to collapse datasets collected on toroids of different sizes into a universal curve on which we perform a global fit. We illustrate the agreement between our final set of parameters and a *d* = *d*(*r*) curve in Fig. 5A.

**Figure 5.**
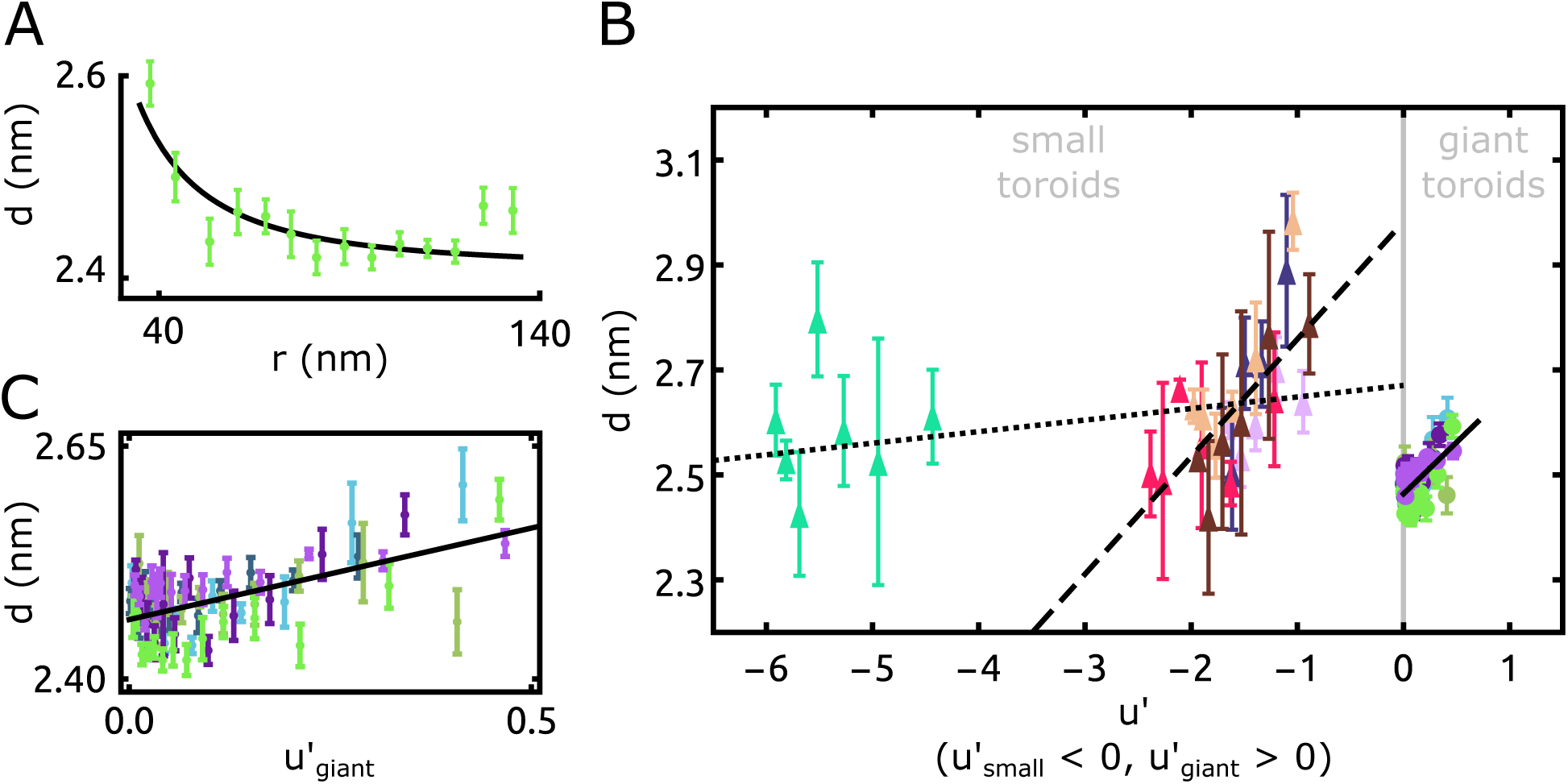
Fitting our model elucidates the dependence of the filament spacing on position and toroid geometry. (A) Example for a single toroid. The line is the fit of the theoretical model 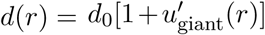, with 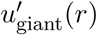 from Eq. (6). The estimated free parameters are *d*_0_ = 2.42 ± 0.01 nm and *ϵ* = 0.12 ± 0.03 (average ± sd). (B) Filament spacing *d* for small (giant) toroids collapses onto a single master curve when expressed as a function of 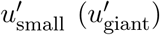. Note that Eq. (5) implies that 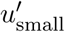 cancels at 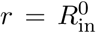, while Eq. (6) implies that 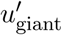 cancels at 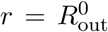. The solid line is the fit to the model of the giant toroid data. The dotted and dashed lines are alternative fits of the data from small toroids. The former includes the leftmost outlier dataset (aquamarine colored triangles), while the latter excludes it. The inferred model’s parameters corresponding to the dashed and the solid lines are provided in Tab. I. Colors and symbols are consistent with Fig. 3. For all datasets, the values of 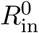 and 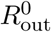 used in the expression of *u*′ correspond to our measurements of the inner and outer radii given in Supplementary Data II. (C) Close-up of the giant toroid data in panel B.

While the curves presented in Fig. 3 are all qualitatively similar, our model predicts that the detailed dependence of the spacing *d* on the radius *r* depends on the toroid-specific parameters *R*_in_ and *R*_out_, preventing a collective fit of all our datasets. Conveniently however, within our small parameter expansion, *d* is an affine function of *u*′(*r*), with parameters *ϵ* and *d*_0_. We thus plot *d* as a function of *u*′ for all our T5 capsid datasets in Fig. 5(B, C). The theory predicts that this dependence should be affine, and we find that our data on small and giant toroids are each separately compatible with this prediction. However the same line clearly cannot be used to fit the two sets, consistent with our earlier comment that the same optimal spacing *d*_0_ cannot be used for both giant and small toroids. To determine the *d*_0_ associated with each dataset, we perform separate least-square fits on small (*u*′ < 0) and giant (*u*′ > 0) toroids, using *ϵ* and *d*_0_ as free parameters, and plot the resulting affine dependences as dotted and solid lines in Fig. 5(B, C). A closer inspection of this figure shows that one of the small toroids is a clear outlier that leads to a large statistical error on the slope of the dotted line (*ϵ* = 0.008 ± 0.004). We thus suggest an alternative fit, excluding it, as a dashed line which we will focus on in our subsequent discussion. We present the inferred values of the free parameters in Tab. I and compute 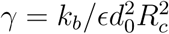 using *R*_*c*_ = 40 nm, *T* = 298 K and 𝓁_*p*_ = 40 nm [which is reasonable under our salt conditions [39]]. Our fits lead to two consistent estimates for the interaction stiffness *γ* with values between 0.17 and 0.21 pN nm^−2^, without statistically meaningful variations going from small to giant toroids. Conversely, the equilibrium spacings *d*_0_ differ significantly between small and giant toroids, shifting from 2.98 to 2.46 nm respectively. This finding suggests that the interactions between DNA helices depend on their curvature, and favor smaller spacings in giant than in small toroids.

**Table I.** Inferred model parameters (average ± sd).

| | $d_0$ (nm) | $\epsilon$ | $\gamma$ (pN nm$^{-2}$) |
| --- | --- | --- | --- |
| Small toroids | $2.98 \pm 0.08$ | $0.07 \pm 0.02$ | 0.17 |
| Giant toroids | $2.46 \pm 0.01$ | $0.08 \pm 0.01$ | 0.21 |

## Discussion

While previous studies on the interactions between DNA helices have largely been focused on straight filaments, our study shows that curving DNA on the scale of a few tens of nanometers modifies these interactions according to two generic mechanisms.

The first mechanism is purely elastic: DNA bending rigidity penalizes the inner regions of toroidal bundles, where helices are more curved, implying that helices are locally depleted from these regions. This mechanism should operate even in the presence of purely repulsive inter-helical forces, *i*.*e*. also in the absence of condensing multivalent cations. This mechanism is indeed reminiscent of the coupling between curvature and density at work in a previous theoretical study [40], where a toroid made of DNA helices interacting through screened Coulomb repulsions is stabilized by a fixed external boundary.

Our second mechanism consists in a hitherto uncharacterized curvature-dependent helix-helix interaction, resulting in a shift of the equilibrium spacing *d*_0_ from 2.46 nm in low-curvature giant toroids to 2.98 nm in high-curvature small ones without a noticeable change in the steepness of the interaction potential. While the microscopic origin of this shift is unclear, we speculate that it could be related to the relative alignment of neighboring DNA helices. Indeed, the inhomogeneous helical charge distribution of phosphate groups and counter-ions causes DNA helices packed at high density to order longitudinally through an “electrostatic zipper” mechanism [41]. Such longitudinal correlations of the helices were characterized using X-ray diffraction patterns [42] and cryo-EM imaging [32]. Consistent with our proposal, these correlations were found to differ in highly curved small toroids and straight bundles [32]. Specifically, while helices in straight DNA bundles were found to align in register (major grooves of one helix facing the minor grooves of its nearest neighbors in all three directions of the hexagonal lattice), alternating longitudinal orientations were found in small confined toroids. We suggest that this mechanism may account for the curvature-dependence reported here, a possibility further supported by previous suggestions that helical correlations may vary with interaxial separation [42, 43], as well as with the lattice geometry [43].

It appears unlikely that the second mechanism could explain the radius-dependent spacing observed in individual toroids without the need to invoke the first. Indeed, helix correlations, unlike their spacing, appear fairly conserved throughout small confined toroids [32]. Nevertheless, a more complete characterization of the variation of helical correlations with curvature would be helpful to better understand this mechanism.

Our mechanisms suggest that the inter-filament spacing in the less curved outer regions of our largest toroids is close to that of a straight DNA bundle. To confirm this prediction, we compare our estimates of the equilibrium inter-helical spacing 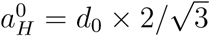 in small 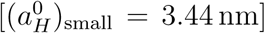 and giant toroids 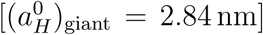 with previous measurements performed in hexagonal packings of short, straight DNA fragments under ionic conditions similar to ours and yielding 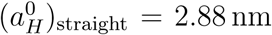 [37]. The virtual indistinguishability of 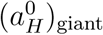 and 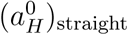 supports our assumption that inter-filament interactions in giant toroids are comparable to those of straight DNA arrays. This should be confirmed by a comprehensive analysis of the relationship between curvature and helical correlations.

Our model also allows us to infer the stiffness per unit DNA length of the locally parabolic interaction potential, with values ranging between 0.17 and 0.21 pN · nm^−2^. These values are rather low compared to those estimated in two previous experimental and numerical works [30, 44]. While the former study presents direct osmotic measurements of these interactions, the latter proposes a microscopic account based on a numerical reconstruction of the cation-mediated force in a hexagonal packing of straight DNA filaments. The authors’ reconstruction of the distance-dependent interfilament force, implies a stiffness of their helix-helix interaction potential close to equilibrium roughly equal to 10 pN·nm^−1^ per helical turn, *i*.*e*., *γ*_straight_ ≃3 pN · nm^−2^, one order of magnitude larger than our estimate. A first factor in explaining this discrepancy could be the ionic conditions considered in these studies, which include spermine and Na^+^ concentrations similar to ours, but not the divalent ions Mg^2+^ and Ca^2+^ present in our experiments. It is furthermore possible that the longitudinal correlations between neighboring helices in highly curved regions could also play a role. Indeed, correlations reported in small confined toroids [32] differ from those found in Ref. [44], which would be interesting to investigate in further numerical simulations.

Overall, our results indicate that curvature tends to increase the spacing between DNA helices in curved assemblies through both DNA elasticity and curvature-dependent helix-helix interactions. The curvature-dependence of the interactions could be related to a modification in their longitudinal correlations, possibly leading to weakened effective interactions. This potential effect of longitudinal correlations may have a measurable impact in various biological contexts. These effects could furthermore be harnessed in the design of curved DNA origami, particularly when curvature is generated by locally tuning the helical longitudinal correlation of tightly packed double helices [4].

## MATERIALS AND METHODS

### Preparation of giant multi-molecular and confined monomolecular toroids

Bacteriohages T5st(0) (Genbank Acc AY692264) and Lambda were obtained by infection of E. coli K12 and E. coli R594 (cI857 S7) strains respectively, purified on cesium chloride gradients, and extensively dialyzed against NaCl 100 mM, MgCl2 1 mM, CaCl2 1 mM, Tris 10 mM pH 7.6. Stock suspensions were stored at 4°C. FhuA and LamB receptors were purified respectively from the strain E. coli AW740 and pop 154 from K12 in which the lamB gene is transduced from Shigella sonnei, as described earlier [45–47], and stock solutions were stored at -80°C. DNA ejection from phages was triggered by mixing the phage stock solution with 110 FhuA/120 LamB molecules per infective phage (T5/Lambda respectively), in the presence of 0.03% LDAO (w:w), and 4 or 40 mM spermine. Bacteriophages were incubated with 4 or 40 mM spermine about 20-30 minutes prior to the addition of the receptor. The ionic environment was kept constant at all stages of the preparations. The phage concentration was adjusted so that the final global DNA concentration *C*_DNA_ = 4 mg/ml. For ejection, the sample was transferred at 37°C during 90 minutes, and transferred back to room temperature before freezing for cryo-EM observation. To obtain multi-molecular giant toroids outside phage capsids, the ejection was simply let to proceed until completion (90 min). To obtain monomolecular confined toroids, DNAse (Invitrogen, 20-25 ue/*µ*l) was added to the sample at t = 5-10 minutes after the addition of FhuA/LamB.

### Cryo Electron Microscopy

3 *µ*l of the toroid suspension were deposited onto a glow-discharged holey carbon grid (Quantifoil R2/2), blotted with a filter paper for 4 seconds, and plunged into liquid ethane cooled down by liquid nitrogen, using a Vitrobot Mark IV (ThermoFisher) operated at room temperature and 100% relative humidity. Frozen samples were transferred and observed in a cryo-TEM, either a JEOL 2010F or a Titan Krios (ThermoFisher) equipped with a Cs corrector, a Quantum GIF energy filter (Gatan, slit width set to 20eV) and a Volta Phase Plate (VPP), operated at 200 kV and 300kV respectively. Images were recorded either at a nominal defocus of 900 nm on Kodak SO163 negative films (JEOL 2010F acquisitions) or close to focus on a Gatan K2 summit camera (Titan Krios with VPP acquisitions). Negative films were developed in full strength Kodak D19 for 12 minutes, and scanned with a Coolscan 9000 (Nikon) at a resolution of 4000 pixels per inch. Images recorded on the JEOL 2010F were denoised by wavelet filtration using ImageJ (“A trous filter” plugin, *k*_1_ = 20, *k*_*n*>1_ = 0). In all cases, micrographs were calibrated using full T5 st(0) phages imaged in the same conditions. The DNA hexagonal lattice spacing in full-filled phages was measured on line profiles, and image scale was calculated from the known value previously recorded by X-ray scattering [48]. Line profiles were recorded using imageJ, with a line width of 5 nm (giant toroids) or 2.65 nm (confined toroids).

## ACKNOWLEDGMENTS

We thank Marta de Frutos (Laboratoire de Physique des Solides, Orsay) for T5 purification, Pascale Boulanger, Madalena Renouard (Institut de Biologie Intégrative de la Cellule, Orsay) and Virginie Bailleux (Laboratoire de Physique des Solides, Orsay) for participation in FhuA and LamB purification. We thank Jéril Degrouard (Laboratoire de Physique des Solides, Orsay) and Julio Ortiz (Institut de Génétique et de Biologie Moléculaire et Cellulaire, Illkirch) for contribution to cryo-EM data acquisition. We thank Emmanuel Trizac and Ivan Palaia (Laboratoire de Physique Théorique et Modèles Statistiques, Orsay) for useful discussions. AL and FL acknowledge support by CNRS, the French Research Agency (ANR-12-BSV5-0023) and Investissements Avenir-LabEx PALM (ANR-10-LABX-0039-PALM), and by the French Infrastructure for Integrated Structural Biology (FRISBI, ANR-10-INBS-05). LB was supported by the “IDI 2016” project funded by the IDEX Paris-Saclay, ANR-11-IDEX-0003-02. ML was supported by Marie Curie Integration Grant PCIG12-GA-2012-334053, “Investissements d’Avenir” LabEx PALM (ANR-10-LABX-0039-PALM), ANR grant ANR-15-CE13-0004-03 and ERC Starting Grant 677532. ML’s group belongs to the CNRS consortium CellTiss.

## Notes

### Competing Interest Statement

The authors have declared no competing interest.

