## Supplementary Material for "Local structure of DNA toroids reveals curvature-dependent intermolecular forces"

#### This PDF includes:

- Supplementary Data I
- Supplementary Data II
- Supplementary Mathematical Modeling

#### Supplementary data I

#### Average lattice spacing measured on top-view cryo-EM images of toroids

| $\bar{d}$ mean ± sd (nm) | 4 mM spermine | | 40 mM spermine | |
|---|---|---|---|---|
| | T5 | λ | T5 | λ |
| Confined toroids | 2.65 ± 0.11 | 2.64 ± 0.08 | 2.64 ± 0.04 | n/a |
| Unconfined toroids | 2.52 ± 0.03 | n/a | 2.51 ± 0.04 | 2.54 ± 0.04 |

### Supplementary data II

#### Experimental conditions, $R_{in}$, $R_{out}$, and averaged d values as a function of R for the different toroids

##### Giant toroids

| | |
|---|---|
| $R_{in}$ (nm) | 37.5 |
| $R_{out}$ (nm) | 135 |
| Number of profiles averaged | 8 |
| Spermine concentration | 40 mM |
| Used bacteriophage species | T5 |

| d (nm) | | R (nm) |
|---|---|---|
| average | $sd/\sqrt{N}$ * | |
| 2.61 | 0.039 | 39.38 |
| 2.57 | 0.043 | 47.20 |
| 2.48 | 0.027 | 55.08 |
| 2.47 | 0.01 | 62.87 |
| 2.49 | 0.016 | 70.32 |
| 2.44 | 0.009 | 77.64 |
| 2.47 | 0.006 | 85.01 |
| 2.44 | 0.011 | 92.36 |
| 2.49 | 0.016 | 99.74 |
| 2.45 | 0.018 | 107.13 |
| 2.48 | 0.008 | 114.56 |
| 2.49 | 0.017 | 121.94 |
| 2.50 | 0.018 | 129.44 |

| | |
|---|---|
| $R_{in}$ (nm) | 42 |
| $R_{out}$ (nm) | 142 |
| Number of profiles averaged | 8 |
| Spermine concentration | 40 mM |
| Used bacteriophage species | T5 |

| d (nm) | | R (nm) |
|---|---|---|
| average | $sd/\sqrt{N}$ * | |
| 2.53 | 0.023 | 49.64 |
| 2.51 | 0.011 | 57.23 |
| 2.51 | 0.016 | 64.75 |
| 2.49 | 0.011 | 72.29 |
| 2.47 | 0.018 | 79.83 |
| 2.49 | 0.010 | 87.25 |
| 2.47 | 0.018 | 94.73 |
| 2.46 | 0.016 | 102.14 |
| 2.48 | 0.018 | 109.66 |
| 2.50 | 0.017 | 117.11 |
| 2.46 | 0.016 | 124.60 |
| 2.48 | 0.014 | 131.98 |
| 2.48 | 0.014 | 139.42 |

| | |
|---|---|
| $R_{in}$ (nm) | 45.5 |
| $R_{out}$ (nm) | 134 |
| Number of profiles averaged | 13 |
| Spermine concentration | 40 mM |
| Used bacteriophage species | T5 |

| d (nm) | | R (nm) |
|---|---|---|
| average | $sd/\sqrt{N}$ * | |
| 2.46 | 0.035 | 47.16 |
| 2.52 | 0.042 | 54.49 |
| 2.51 | 0.016 | 62.07 |
| 2.48 | 0.023 | 69.60 |
| 2.47 | 0.021 | 77.04 |
| 2.49 | 0.010 | 84.45 |
| 2.47 | 0.014 | 91.86 |
| 2.49 | 0.023 | 99.31 |
| 2.46 | 0.021 | 106.78 |
| 2.48 | 0.023 | 114.15 |
| 2.52 | 0.030 | 121.63 |

| | |
|---|---|
| $R_{in}$ (nm) | 40 |
| $R_{out}$ (nm) | 140 |
| Number of profiles averaged | 8 |
| Spermine concentration | 40 |
| Used bacteriophage species | T5 |

| d (nm) | | R (nm) |
|---|---|---|
| average | $sd/\sqrt{N}$ * | |
| 2.58 | 45.67 | 0.0215 |
| 2.53 | 53.38 | 0.0268 |
| 2.49 | 60.91 | 0.0257 |
| 2.47 | 68.44 | 0.0265 |
| 2.43 | 75.87 | 0.0151 |
| 2.51 | 83.22 | 0.0216 |
| 2.45 | 90.68 | 0.0177 |
| 2.48 | 98.08 | 0.0554 |
| 2.44 | 105.56 | 0.0182 |
| 2.46 | 112.77 | 0.0234 |
| 2.49 | 120.17 | 0.0204 |
| 2.52 | 127.70 | 0.0102 |

* Note that the number of measures may vary between profiles in a toroid. Standard deviation is not available where only a single measure is possible.

| | |
|---|---|
| $R_{in}$ (nm) | 37.5 |
| $R_{out}$ (nm) | 142 |
| Number of profiles averaged | 10 |
| Spermine concentration | 40 mM |
| Used bacteriophage species | T5 |

| d (nm) | | R (nm) |
|---|---|---|
| average | $sd/\sqrt{N}$ * | |
| 2.59 | 0.022 | 37.68 |
| 2.50 | 0.024 | 44.33 |
| 2.44 | 0.023 | 53.32 |
| 2.47 | 0.022 | 60.70 |
| 2.46 | 0.017 | 68.02 |
| 2.44 | 0.023 | 74.67 |
| 2.42 | 0.017 | 81.39 |
| 2.43 | 0.017 | 88.69 |
| 2.42 | 0.012 | 95.92 |
| 2.43 | 0.012 | 103.23 |
| 2.43 | 0.009 | 110.51 |
| 2.43 | 0.011 | 117.79 |
| 2.47 | 0.018 | 125.20 |
| 2.47 | 0.022 | 132.95 |

| | |
|---|---|
| $R_{in}$ (nm) | 30 |
| $R_{out}$ (nm) | 140 |
| Number of profiles averaged | 6 |
| Spermine concentration | 40 mM |
| Used bacteriophage species | λ |

| d (nm) | | R (nm) |
|---|---|---|
| average | $sd/\sqrt{N}$ * | |
| 2.76 | 0.031 | 34.75 |
| 2.64 | 0.005 | 43.03 |
| 2.54 | 0.005 | 50.94 |
| 2.57 | 0.028 | 58.54 |
| 2.39 | 0.026 | 66.25 |
| 2.53 | 0.038 | 73.42 |
| 2.49 | 0.060 | 81.02 |
| 2.60 | 0.008 | 89.05 |
| 2.49 | 0.031 | 96.84 |
| 2.53 | 0.048 | 104.32 |

| | |
|---|---|
| $R_{in}$ (nm) | 39.5 |
| $R_{out}$ (nm) | 140 |
| Number of profiles averaged | 8 |
| Spermine concentration | 4 mM |
| Used bacteriophage species | T5 |

| d (nm) | | R (nm) |
|---|---|---|
| average | $sd/\sqrt{N}$ * | |
| 2.55 | 0.014 | 39.21 |
| 2.53 | 0.010 | 46.85 |
| 2.53 | 0.005 | 54.43 |
| 2.50 | 0.012 | 62.03 |
| 2.51 | 0.015 | 69.55 |
| 2.50 | 0.016 | 77.09 |
| 2.49 | 0.016 | 84.58 |
| 2.49 | 0.019 | 92.06 |
| 2.50 | 0.015 | 99.54 |
| 2.50 | 0.017 | 107.08 |
| 2.46 | 0.007 | 114.60 |
| 2.49 | 0.009 | 121.95 |
| 2.50 | 0.011 | 129.33 |
| 2.49 | 0.023 | 102.63 |

* Note that the number of measures may vary between profiles in a toroid. Standard deviation is not available where only a single measure is possible.

#### Small confined toroids

| | |
|---|---|
| $R_{in}$ (nm) | 16 |
| $R_{out}$ (nm) | 36 |
| Number of profiles averaged | 5 |
| Spermine concentration | 40 mM |
| Used bacteriophage species | T5 |

| d (nm) | | R (nm) |
|---|---|---|
| average | $sd/\sqrt{N}$ * | |
| 2.98 | 0.055 | 20.85 |
| 2.72 | 0.106 | 23.83 |
| 2.63 | 0.028 | 26.55 |
| 2.56 | 0.061 | 29.18 |
| 2.61 | 0.052 | 31.74 |
| 2.63 | 0.032 | 34.13 |

| | |
|---|---|
| $R_{in}$ (nm) | 16 |
| $R_{out}$ (nm) | 36 |
| Number of profiles averaged | 5 |
| Spermine concentration | 40 mM |
| Used bacteriophage species | T5 |

| d (nm) | | R (nm) |
|---|---|---|
| average | $sd/\sqrt{N}$ * | |
| 2.79 | 0.042 | 19.86 |
| 2.77 | 0.088 | 22.64 |
| 2.60 | 0.095 | 25.41 |
| 2.56 | 0.074 | 28.01 |
| 2.42 | 0.084 | 30.55 |
| 2.53 | - | 32.98 |
| 2.59 | - | 35.66 |

| | |
|---|---|
| $R_{in}$ (nm) | 10 |
| $R_{out}$ (nm) | 36 |
| Number of profiles averaged | 4 |
| Spermine concentration | 4 mM |
| Used bacteriophage species | T5 |

| d (nm) | | R (nm) |
|---|---|---|
| average | $sd/\sqrt{N}$ * | |
| 2.84 | - | 14.89 |
| 2.61 | 0.089 | 17.80 |
| 2.52 | 0.235 | 20.56 |
| 2.58 | 0.105 | 23.16 |
| 2.80 | 0.109 | 25.97 |
| 2.43 | 0.118 | 28.64 |
| 2.53 | 0.037 | 31.20 |
| 2.60 | 0.067 | 33.75 |

| | |
|---|---|
| $R_{in}$ (nm) | 17 |
| $R_{out}$ (nm) | 36 |
| Number of profiles averaged | 6 |
| Spermine concentration | 4 mM |
| Used bacteriophage species | T5 |

| d (nm) | | R (nm) |
|---|---|---|
| average | $sd/\sqrt{N}$ * | |
| 2.89 | 0.145 | 23.83 |
| 2.71 | 0.081 | 26.72 |
| 2.71 | 0.087 | 29.43 |
| 2.50 | 0.108 | 32.14 |
| 2.56 | 0.067 | 34.64 |

* Note that the number of measures may vary between profiles in a toroid. Standard deviation is not available where only a single measure is possible.

| | |
|---|---|
| $R_{in}$ (nm) | 17 |
| $R_{out}$ (nm) | 36 |
| Number of profiles averaged | 10 |
| Spermine concentration | 4 mM |
| Used bacteriophage species | T5 |

| d (nm) | | R (nm) |
|---|---|---|
| average | $sd/\sqrt{N}$ * | |
| 2.64 | 0.155 | 22.34 |
| 2.70 | 0.171 | 24.96 |
| 2.59 | 0.143 | 27.64 |
| 2.53 | 0.179 | 30.24 |
| 2.60 | 0.109 | 32.83 |

| | |
|---|---|
| $R_{in}$ (nm) | 15 |
| $R_{out}$ (nm) | 37 |
| Number of profiles averaged | 3 |
| Spermine concentration | 4 mM |
| Used bacteriophage species | T5 |

| d (nm) | | R (nm) |
|---|---|---|
| average | $sd/\sqrt{N}$ * | |
| 2.64 | 0.127 | 19.36 |
| 2.48 | 0.042 | 22.00 |
| 2.56 | 0.158 | 24.49 |
| 2.66 | 0.018 | 27.04 |
| 2.49 | 0.187 | 29.71 |
| 2.50 | 0.081 | 32.20 |
| 2.58 | - | 34.79 |

| | |
|---|---|
| $R_{in}$ (nm) | 13.5 |
| $R_{out}$ (nm) | 26 |
| Number of profiles averaged | 4 |
| Spermine concentration | 4 mM |
| Used bacteriophage species | λ |

| d (nm) | | R (nm) |
|---|---|---|
| average | $sd/\sqrt{N}$ * | |
| 2.70 | 16.88 | 0.096 |
| 2.50 | 19.58 | 0.077 |
| 2.53 | 22.08 | 0.076 |

* Note that the number of measures may vary between profiles in a toroid. Standard deviation is not available where only a single measure is possible.

### Supplementary Mathematical Modeling

In this supplement, we show how to minimize the free energy of Eq. (3) in the main text following a perturbative approach. The procedure is identical for confined and unconfined toroids, with the only difference that the free and fixed boundaries are inverted in the two cases. Here we specialize to the case of giant toroids, which have fixed internal radius $R_{in} = R_c$ and free outer radius $R_{out}$, such that their free energy is

$$\mathcal{F}[u(r), R_{out}] = \int_{R_c}^{R_{out}} \frac{dr}{d_0} 2\pi[r + \epsilon u(r)] \left\{ \frac{k_b}{2} \frac{1}{[r + \epsilon u(r)]^2} + \frac{\gamma}{2} \left[\epsilon u'(r)\right]^2 - \mu \right\}. \tag{1}$$

Notice that the integral is performed over the *reference state* (see main text).

We proceed as follows. We start by solving force balance equations

$$\left. \frac{\delta \mathcal{F}\left[u(r), R_{\text{out}}\right]}{\delta u(r)} \right|_{R_{\text{out}},\ \mu} = 0, \tag{2a}$$

$$\left. \frac{\partial \mathcal{F}\left[u(r), R_{\text{out}}\right]}{\partial R_{\text{out}}} \right|_{u(r),\ \mu} = 0, \tag{2b}$$

to leading order in the small parameter $\epsilon$. These are solved for expressions of $u(r)$ and $R_{\text{out}}$ in terms of the Lagrange multiplier $\mu$, which we then plug into the constraint equation [Eq. (4) in the main text]

$$\mathcal{L} = \int_{R_c}^{R_{out}} \frac{dr}{d_0} 2\pi[r + \epsilon u(r)] \tag{3}$$

so as to find the dependency of $\mu$ on the input parameters $R_{\text{c}}$ and $\mathcal{L}$.

To develop the perturbative expansion in powers of $\epsilon$, we introduce the non-dimensional units $R_{\text{c}} = \gamma d_0^2 = 1$, yielding the rescaled quantities

$$\tilde{\mathcal{F}} = \frac{\mathcal{F}}{\gamma d_0^2 R_{\text{c}}}, \quad \tilde{R}_{\text{out}} = \frac{R_{\text{out}}}{R_{\text{c}}}, \quad \tilde{r} = \frac{r}{R_{\text{c}}}, \quad \tilde{d}_0 = \frac{d_0}{R_{\text{c}}}, \quad \tilde{u} = \frac{u}{R_{\text{c}}}, \quad \tilde{k}_b = \epsilon \quad \text{and} \quad \tilde{\mu} = \frac{\mu}{\gamma d_0^2}. \tag{4}$$

In these units, Eq. (1) reads:

$$\tilde{\mathcal{F}}\left[\tilde{u}\left(\tilde{r}\right), \tilde{R}_{\text{out}}\right] = \int_1^{\tilde{R}_{\text{out}}} \frac{\mathrm{d}\tilde{r}}{\tilde{d}_0} 2\pi \left[\tilde{r} + \epsilon \tilde{u}\left(\tilde{r}\right)\right] \left\{ \frac{\epsilon}{2} \frac{1}{\left[\tilde{r} + \epsilon \tilde{u}\left(\tilde{r}\right)\right]^2} + \frac{1}{2} \left[\epsilon \tilde{u}'\left(\tilde{r}\right)\right]^2 - \tilde{\mu} \right\}. \tag{5}$$

We can now expand Eq. (2a) in powers of $\epsilon$ and solve it to leading order. Henceforth, we assume that $\mathcal{O}(\tilde{\mu}) = \mathcal{O}(\epsilon)$, given that the total DNA length is automatically conserved in the absence of deformation (*i.e.*, when $\epsilon = 0$). We expand the free energy

[Eq. (5)] as $\tilde{\mathcal{F}} = \tilde{\mathcal{F}}_1\epsilon + \tilde{\mathcal{F}}_2\epsilon^2 + \mathcal{O}(\epsilon^3)$, with:

$$\tilde{\mathcal{F}}_1\left(\tilde{R}_{\text{out}}\right) = \int_1^{\tilde{R}_{\text{out}}} \mathrm{d}\tilde{r}\ 2\pi\left(\frac{1}{2\tilde{r}} - \tilde{\mu}\tilde{r}\right); \tag{6a}$$

$$\tilde{\mathcal{F}}_2\left[\tilde{u}(\tilde{r}), \tilde{R}_{\text{out}}\right] = \int_1^{\tilde{R}_{\text{out}}} \mathrm{d}\tilde{r}\ 2\pi\left\{-\frac{\tilde{u}(\tilde{r})}{2\tilde{r}^2} + \frac{\tilde{r}}{2}\left[\tilde{u}'(\tilde{r})\right]^2 - \tilde{\mu}\tilde{u}(\tilde{r})\right\}. \tag{6b}$$

The leading order of Eq. (2a) is then $\delta\tilde{\mathcal{F}}_2/\delta\tilde{u}(\tilde{r}) = 0$. To calculate the variational derivative, we introduce a small perturbation $\delta\tilde{u}(\tilde{r})$ in the functional $\tilde{\mathcal{F}}_2$, getting

$$\tilde{\mathcal{F}}_2[\tilde{u}(\tilde{r}) + \delta\tilde{u}(\tilde{r}), R_{\text{out}}] = \tilde{\mathcal{F}}_2[\tilde{u}(\tilde{r}), R_{\text{out}}] + \delta\tilde{\mathcal{F}}_2[\tilde{u}(\tilde{r}), \delta\tilde{u}(\tilde{r}), R_{\text{out}}] + \mathcal{O}\left[\delta\tilde{u}(\tilde{r})^2\right], \tag{7}$$

where

$$\delta\tilde{\mathcal{F}}_2\left[\tilde{u}(\tilde{r}), \delta\tilde{u}(\tilde{r}), R_{\text{out}}\right] = \int_1^{\tilde{R}_{\text{out}}} \mathrm{d}\tilde{r}'\ 2\pi\left[-\frac{\delta\tilde{u}(\tilde{r})}{2\tilde{r}^2} + \tilde{r}\tilde{u}'(\tilde{r})\,\delta\tilde{u}'(\tilde{r}) - \tilde{\mu}\delta\tilde{u}(\tilde{r})\right]. \tag{8}$$

Integrating by parts the second term in Eq. (8), we obtain

$$\delta\tilde{\mathcal{F}}_2\left[\tilde{u}(\tilde{r}), \delta\tilde{u}(\tilde{r}), R_{\text{out}}\right] = \left[\tilde{r}\tilde{u}'(\tilde{r})\,\delta\tilde{u}(\tilde{r})\right]_1^{\tilde{R}_{\text{out}}} + \int_1^{\tilde{R}_{\text{out}}} \mathrm{d}\tilde{r}'\ 2\pi\left\{-\frac{1}{2\tilde{r}^2} - \left[\tilde{r}\tilde{u}'(\tilde{r})\right]' - \tilde{\mu}\right\}\delta\tilde{u}(\tilde{r}). \tag{9}$$

Hence, the stationarity condition $\delta\tilde{\mathcal{F}}_2/\delta\tilde{u}(\tilde{r}) = 0$ boils down to a differential equation,

$$\frac{1}{2\tilde{r}^2} + \left[\tilde{r}\tilde{u}'(\tilde{r})\right]' + \tilde{\mu} = 0, \tag{10}$$

and two boundary conditions,

$$\tilde{u}(1) = 0, \tag{11a}$$

$$\tilde{u}'(\tilde{R}_{\text{out}}) = 0. \tag{11b}$$

Eq. (11a) is a fixed boundary condition indicating that the innermost DNA filament sticks to the viral capsid, while Eq. (11b) is a free boundary condition indicating that the toroid is unconfined. The solution of Eqs. (10) and (11) is

$$\tilde{u}(\tilde{r}) = \frac{(\tilde{r}-1)(1-2\tilde{r}\tilde{\mu})}{2\tilde{r}} - \left(\frac{1}{2\tilde{R}_{\text{out}}} - \tilde{\mu}\tilde{R}_{\text{out}}\right)\log(\tilde{r}). \tag{12}$$

Turning to Eq. (2b), we call $f(r, u(r), u'(r))$ the integrand of Eq. (5) and rewrite the left hand side of Eq. (2b) as

$$\frac{\partial}{\partial\tilde{R}_{\text{out}}}\int_1^{\tilde{R}_{\text{out}}} \mathrm{d}\tilde{r}\ \tilde{f}(\tilde{r}, \tilde{u}(\tilde{r}), \tilde{u}'(\tilde{r})) = \tilde{f}(\tilde{R}_{\text{out}}, \tilde{u}(\tilde{R}_{\text{out}}), \tilde{u}'(\tilde{R}_{\text{out}})). \tag{13}$$

Combining the equation above with Eq. (11b), Eq. (2b) becomes

$$\tilde{f}(\tilde{R}_{\text{out}}, \tilde{u}(\tilde{R}_{\text{out}})) = 0. \tag{14}$$

We solve this equation perturbatively, this time expanding the outer radius in powers of $\epsilon$: $\tilde{R}_{\text{out}} = \tilde{R}^0_{\text{out}} + \mathcal{O}(\epsilon)$. To first order in $\epsilon$, Eq. (14) reads $1/(2\tilde{R}^0_{\text{out}}) - \tilde{\mu}\tilde{R}^0_{\text{out}} = 0$, yielding

$$\tilde{R}^0_{\text{out}} = \frac{1}{\sqrt{2\tilde{\mu}}}. \tag{15}$$

We now solve the constraint equation [Eq. (3)] for $\mu$. The zeroth order in $\epsilon$ of the equation is

$$\tilde{\mathcal{L}} = \int_1^{\tilde{R}^0_{\text{out}}} \frac{\mathrm{d}\tilde{r}}{\tilde{d}_0} \tilde{r}, \tag{16}$$

which gives $\tilde{\mu} = [2(\tilde{d}_0\tilde{\mathcal{L}}/\pi+1)]^{-1}$. Plugging the expression of $\tilde{\mu}$ into Eqs. (12) and (15), we get the displacement field and the outer radius in terms of the input parameters:

$$\tilde{u}\,(\tilde{r}) = \frac{1}{2\tilde{r}}\,(\tilde{r}-1)\left[1 - \frac{\tilde{r}}{\left(\tilde{R}^0_{\text{out}}\right)^2}\right]; \tag{17a}$$

$$\tilde{R}^0_{\text{out}} = \sqrt{\tilde{d}_0\tilde{\mathcal{L}}/\pi + 1}. \tag{17b}$$

Going back to dimensional units and taking the first derivative of Eq. (17a), we finally get Eq. (6) of the main text.
